# Genome assembly of a diversity panel of *Chenopodium quinoa*

**DOI:** 10.1101/2024.07.07.602379

**Authors:** Elodie Rey, Michael Abrouk, Isabelle Dufau, Nathalie Rodde, Noha Saber, Jana Cizkova, Gabriele Fiene, Clara Stanschewski, David E Jarvis, Eric N Jellen, Peter J Maughan, Ingrid von Baer, Maxim Troukhan, Maksym Kravchuk, Eva Hribova, Stephane Cauet, Simon G. Krattinger, Mark Tester

## Abstract

Quinoa (*Chenopodium quinoa*) is an important crop for the future challenges of food and nutrient security. Deep characterization of quinoa diversity is needed to support the agronomic improvement and adaptation of quinoa as its worldwide cultivation expands. In this study, we report the construction of chromosome-scale genome assemblies of eight quinoa accessions covering the range of phenotypic and genetic diversity of both lowland and highland quinoas. The assemblies were produced from a combination of PacBio HiFi reads and Bionano Saphyr optical maps, with total assembly sizes averaging 1.28 Gb with a mean N50 of 71.1 Mb. Between 43,733 and 48,564 gene models were predicted for the eight new quinoa genomes, and on average, 66% of each quinoa genome was classified as repetitive sequences. Alignment between the eight genome assemblies allowed the identification of structural rearrangements including inversions, translocations, and duplications. These eight novel quinoa genome assemblies provide a resource for association genetics, comparative genomics, and pan-genome analyses for the discovery of genetic components and variations underlying agriculturally important traits.

## Background & Summary

Over the past fifty years, quinoa (*Chenopodium quinoa* Willd.) cultivation transformed from the state of a subsistence crop for local farmers exclusively grown in the Andean region of South America, to being cultivated or experimented with by more than 110 countries across all continents^1^. The rapid global expansion of quinoa beyond the Andes arose from the international recognition of the high nutritional value of its seeds, resilience to environmental extremes, tolerance to several abiotic stresses, and the large genetic diversity available for this crop. These qualities make quinoa a promising future crop, specifically for diversifying sustainable cropping systems and food markets which in turn will address nutrient and food security in marginal agricultural regions of the world^2–4^. Along with the spread of quinoa cultivation comes the need to develop breeding programs, collect and catalog the available genetic diversity, and identify favorable allelic variation underlying beneficial traits that will allow quinoa to adapt to new environments and agronomic practices^5,6^. Underlying these breeding program and genetics studies is the need to generate genomic resources specific to quinoa to support its accelerated improvement.

Quinoa germplasm banks are composed of several thousand accessions, both landraces and cultivars, fromdiverse geographical origins^7,8^. Diversity studies based on molecular markers classified quinoa into two major groups: 1) highland quinoa, which displays the greatest genetic diversity and whichspans the high plains of Bolivia, Peru, and Ecuador near its center of origin by Lake Titicaca (between 3650 and 4200 m above sea level); and 2) lowland quinoa, also named coastal or sea-level quinoa, which defines a unique South-Central Chilean ecotype of lesser diversity and has been used to breed quinoa for lowland regions worldwide^2,8,9^. Additional studies grouped quinoa according to morphological traits linked with adaptation to specific agro-ecological conditions in major production areas, further classifying quinoa into four smaller genetic groups also referred to as ecotypes (Valley, Northern and Southern Highland, and Sea Level)^10–14^. Phenotypic diversity and genetic variation for adaptation to various environments and stresses have been described within each regional geographical distribution. However, very little is known of the natural allelic variation underlying these traits since genomic resources are still restricted to a limited representation of the available genetic diversity.

Several assemblies have been produced for the quinoa genome (2*n* = 4*x* = 36) in the past decade and have been important for these initial characterizations of quinoa genetic diversity and genetic studies. Draft assemblies have been produced for four quinoa accessions, including Kd^15^, a Japanese inbred accession; PI 614886 (QQ74)^16^, a coastal Chilean accession; Real^17^, a highland accession representing the most commonly cultivated commercial variety; and the Bolivian accession CHEN125^18^. Though fragmented and incomplete, these resources have enabled transcriptomic analysis and targeted gene family approaches for the identification and mapping of genes involved in adaptation and domestication traits including seed saponin synthesis^16^, quinoa seed quality^19^, flowering time^20^, and the response to abiotic^17,21,22–25^ and biotic^26^ stresses. Recently, we produced an improved version, both in terms of contiguity and completeness, of the coastal quinoa PI 614886 genome assembly (QQ74-V2^27^) which has enabled larger-scale genetic studies^28–33^.(both in terms of population size and allelic variation representation) through quantitative genetics and genome-wide association studies. This new assembly, together with the resequencing of a set of quinoa accessions from both highland and lowland groups, uncovered large genomic rearrangements which may have significant implications for quinoa breeding and improvement. The results of that study further emphasized the need to consider not only nucleotide but also structural polymorphism to better understand quinoa diversity, especially as structural variants can be important components underlying phenotypic variation.

Here we present the genome assembly and annotation of a panel of eight quinoa accessions of diverse geographical origins selected to represent the diversity of quinoa phenotypes and to support the genetic improvement and adaptation of quinoa to warm and arid environments. All eight genomes are high-quality, chromosome-scale reference sequences assembled from over 30x genome coverage of PacBio HiFi long-reads, with validation and scaffolding provided by Bionano Saphyr optical maps. We further provide a map of structural rearrangements (inversion, duplication, and translocation) between the eight genomes. These genomes represent important resources for the investigation of the genetic components underlying important agronomic traits for quinoa improvement needed to meet the challenges of adapting to novel environments outside its native cultivation range.

## Methods

### Plant material

A set of eight quinoa accessions, including three lowland (Regalona and Javi from Chile; D-12282 from Argentina) and five highland genotypes (CHEN-199 and PI-614919 from Bolivia; 03-21-072RM, D-10126 and CHEN-90 from Peru), were primarily selected to represent the diversity of phenotypes (growth habit, panicle architecture, leaf shape, stem and seed colors – Fig. 1) and within the selected diversity for performing relatively well in hot, dry and short-day environments^34^. Each of these eight accessions is highly homogeneous and exhibits low seed shattering^34^. Previous analysis of SNPs produced through short-read resequencing indicated that all eight genotypes have high coefficient of inbreeding (F.coef ≥ 0.85; homozygous) and are well distributed across the spread of genetic diversity of quinoa^35^.

**Fig. 1:**
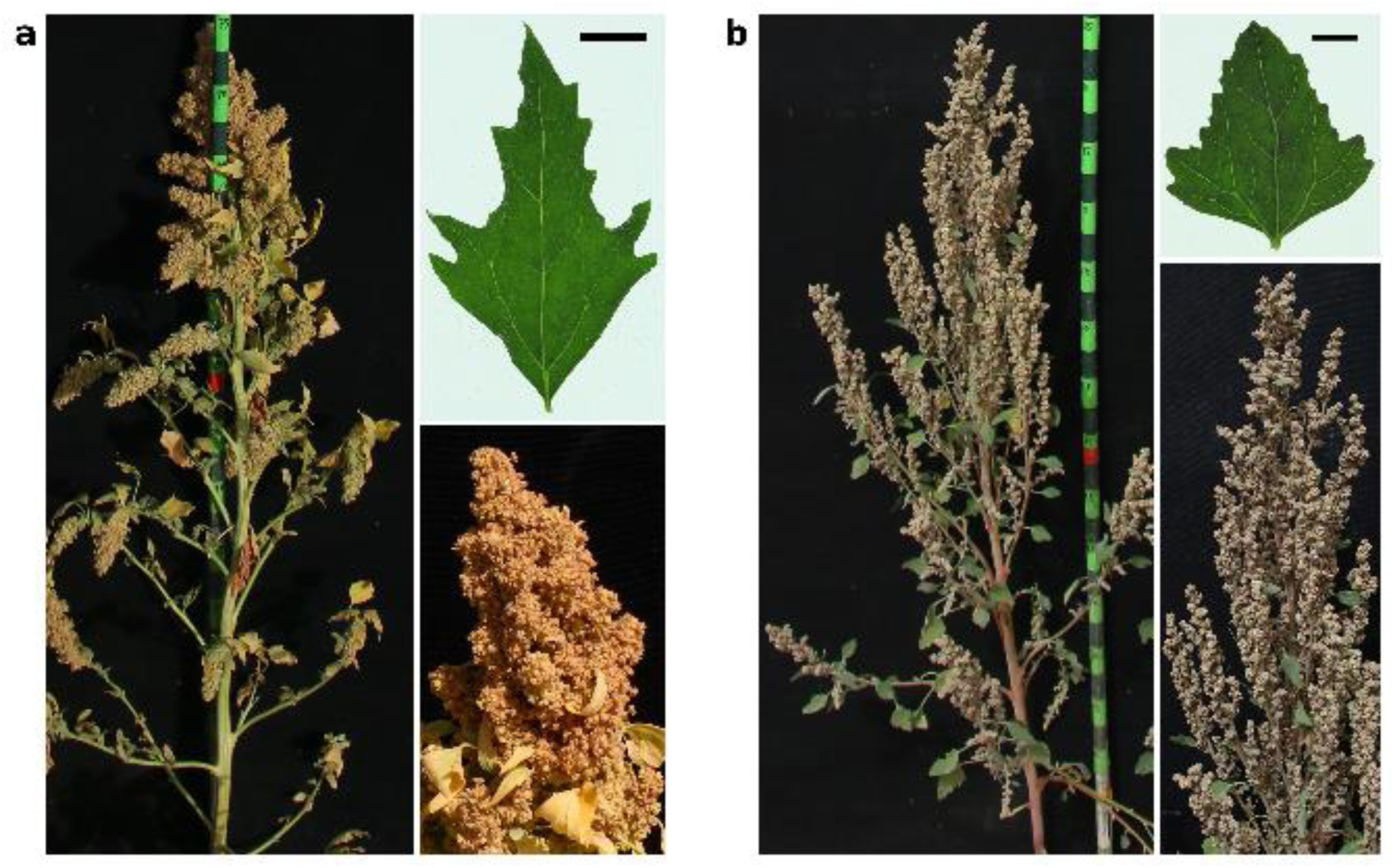
Phenotypic diversity between highland and lowland quinoas. Photos depicting the contrasting phenotypes for plant architecture, leaf shape and panicle between (a) the lowland accession Regalona and (b) the highland accession 03-21-072RM. Photos are from 2019 field trials in Hada Al-Sham, Saudi Arabia.

### Reference genome sequencing and assembly

#### Genome size estimation

We performed genome size estimation for each of the eight quinoa genotypes to inform the genome sequencing and assembly strategy. DNA amounts of each *Chenopodium quinoa* accession were estimated using CytoFLEX flow cytometer (Beckman Coulter, Brea, USA) equipped with a 488 nm solid-state laser. Samples were prepared following the protocol of Doležel et al.^36^ using LB01 nuclei isolation buffer and propidium iodide staining. *Lycopersicon esculentum* cv. Stupicke polni tyckove rane (2C = 1.96 pg) was used as the internal reference standard^37^. Five individuals were analyzed per accession, and each plant was analyzed three times on three different days. 2C nuclear DNA amounts were calculated according to Doležel et al.^36^. Genome sizes (1C values) were then determined considering 1 pg DNA equal to 0.978×10^9^ bp according to Doležel et al ^38^. The genome sizes averaged 1.42 Gb with limited variation between accessions, which is in agreement with previous studies^39,40^. The smallest genome (1.40 Gb) was found for CHEN-199 and the largest (1.43 Gb) for Javi, representing less than 2% genome size difference (Table 1).

**Table 1.** Genome size estimation.

| **Genotype** | **2C nuclear DNA content [pg]** |  | **Genome size**<br>**[Gbp/1C]** |
| --- | --- | --- | --- |
|  | **Mean** | **± SD** |  |
| D-12282 | 2.929 | 0.035 | 1.432 |
| Regalona | 2.883 | 0.029 | 1.41 |
| Javi | 2.932 | 0.033 | 1.434 |
| D-10126 | 2.885 | 0.066 | 1.411 |
| PI-614919 | 2.906 | 0.04 | 1.421 |
| CHEN-90 | 2.925 | 0.025 | 1.43 |
| 03-21-072RM | 2.928 | 0.023 | 1.432 |
| CHEN-199 | 2.879 | 0.023 | 1.408 |

#### DNA extraction and sequencing

The seed stock used for genome sequencing is the second generation of propagation through single-seed descent from the original seeds received from the genebanks. Seeds were sown in pots containing a mixture of 30% sand + 70% soil & rocks (3:1 parts), and grown in KAUST greenhouses in short-day conditions (10-hour day length) with 26°C day / 18°C night temperatures. Quinoa plants at the bolting stage (50 on the Biologische Bundesanstalt Bundessortenamt und CHemische Industrie (BBCH)-scale ^41,42^) were dark-treated for 48 hours, at which point the youngest leaf material from a single plant for each genotype was collected and immediately frozen in liquid nitrogen and kept at −80°C until DNA extraction. Samples were ground into powder in liquid nitrogen using mortar and pestle, and high molecular weight (HMW) genomic DNA was isolated using the LeafGo protocol^43^ for long-read sequencing. DNA was quantified using a Qubit dsDNA HS Assay (Q32851, ThermoFisher Scientific), purity was confirmed using a Nanodrop spectrophotometer by checking that 260/280 and 260/230 ratios and DNA fragments size (>50 kb) was validated using the FemtoPulse system (Agilent, Santa Clara, CA, USA). HiFi libraries were then prepared according to the manual “Procedure & Checklist - Preparing HiFi SMRTbell® Libraries using the SMRTbell Express Template Prep Kit 2.0” (PN 101-853-100, Pacific Biosciences, Menlo Park, CA, USA) with 10 μg DNA sheared using the Megaruptor 2 system (Diagenode, Liège Science Park, Belgium) to obtain an average fragment size of 15-20 kb. Size-selected libraries were sequenced on a PacBio Sequel II system in CCS mode for 30 hours. For each accession, two SMRT cells were sequenced, and an average of 53 Gb PacBio HiFi reads per sample were obtained, corresponding to a ∼36-fold coverage (Table 2).

**Table 2:** Genome sequencing and assembly statistics.

| **Genotype** | **03-21-072RM** | **CHEN-199** | **CHEN-90** | **D-10126** | **D-12282** | **Javi** | **PI-614919** | **Regalo na** |
| --- | --- | --- | --- | --- | --- | --- | --- | --- |
| **HiFi reads** |  |  |  |  |  |  |  |  |
| Mean read quality | 31.35 | 32.35 | 33.40 | 31.95 | 31.45 | 30.15 | 32.80 | 31.25 |
| Expected coverage (x) | 41.56 | 37.47 | 29.19 | 44.2 | 34.01 | 31.94 | 35.96 | 40.47 |
| **Contig Assembly** |  |  |  |  |  |  |  |  |
| Number of contigs | 692 | 916 | 2,201 | 540 | 693 | 554 | 919 | 701 |
| Total assembly length (bp) | 1,311,128,858 | 1,306,739,747 | 1,881,510,923 | 1,291,635,203 | 1,305,091,460 | 1,313,180,962 | 1,302,672,302 | 1,318,261,280 |
| N50 | 64,501,058 | 43,735,722 | 9,804,934 | 24,917,000 | 25,043,658 | 61,012,816 | 30,750,493 | 45,520,693 |
| N90 | 19,882,343 | 13,039,303 | 423,497 | 7,272,617 | 9,383,707 | 21,330,726 | 9,603,103 | 15,281,957 |
| **Hybrid scaffold** |  |  |  |  |  |  |  |  |
| Number of contigs | 21 | 21 | 27 | 22 | 20 | 24 | 22 | 28 |
| Total assembly length (bp) | 1,283,791,981 | 1,274,953,502 | 1,283,209,458 | 1,275,934,985 | 1,280,906,559 | 1,288,166,338 | 1,297,375,796 | 1,289,500,925 |
| N50 | 71,031,185 | 70,960,791 | 71,021,482 | 71,110,407 | 71,728,093 | 71,904,790 | 71,076,969 | 70,040,807 |
| N90 | 59,961,463 | 59,226,503 | 40,453,756 | 63,317,150 | 60,203,356 | 60,571,168 | 59,411,264 | 37,849,831 |
| **Not scaffolded** |  |  |  |  |  |  |  |  |
| Number of contigs | 650 | 862 | 2,107 | 448 | 621 | 516 | 852 | 649 |
| Total assembly length (bp) | 27,503,412 | 31,988,790 | 601,040,957 | 18,670,849 | 24,606,198 | 25,069,896 | 33,146,976 | 29,242,991 |
| N50 | 40,049 | 35,375 | 658,713 | 40,635 | 38,865 | 48,830 | 36,054 | 43,583 |
| N90 | 29,973 | 24,927 | 154,646 | 28,869 | 27,842 | 32,995 | 26,203 | 30,487 |
| **BUSCO (embryophyta_odb10)** |  |  |  |  |  |  |  |  |
| Complete (%) | 98 | 98 | 981 | 98.1 | 98.1 | 98.1 | 98 | 98 |
| Single-copy (%) | 9.5 | 9.9 | 9.8 | 9.8 | 9.9 | 9.8 | 9.8 | 9.5 |
| Duplicated (%) | 88.5 | 88.1 | 88.3 | 88.3 | 88.2 | 88.3 | 88.2 | 88.5 |
| Fragmented (%) | 0.2 | 0.2 | 0.002 | 0.2 | 0.2 | 0.2 | 0.2 | 0.2 |
| Missing (%) | 1.8 | 1.8 | 0.017 | 1.7 | 1.7 | 1.7 | 1.8 | 1.8 |
| **Merqury** |  |  |  |  |  |  |  |  |
| Consensus<br>quality value<br>(QV) | 66.22 | 64.72 | 67.04 | 66.60 | 65.69 | 65.42 | 69.03 | 68.87 |
| K-mer<br>completeness<br>(%) | 98.93 | 98.78 | 98.39 | 99.03 | 98.98 | 98.41 | 99.00 | 98.25 |

**Table 3:** Repeat element annotation.

| Genot<br>ype | 03-21-072R<br>M | CHEN<br>-199 | CHEN<br>-90 | D-10126 | D-12282 | Javi | PI-614919 | Regalo<br>na | Averag<br>e |
| --- | --- | --- | --- | --- | --- | --- | --- | --- | --- |
| Class |  |  |  |  |  |  |  |  |  |
| LTR |  |  |  |  |  |  |  |  |  |
| Copia | 7.89% | 8.53% | 8.06% | 7.93% | 8.04% | 7.77% | 8.65% | 8.42% | **8.16%** |
| Gypsy | 29.21% | 30.25% | 35.25% | 30.84% | 35.17% | 28.41% | 36.76% | 35.07% | **32.62%** |
| unkno<br>wn | 10.91% | 9.23% | 10.39% | 9.71% | 9.91% | 11.40% | 9.21% | 10.72% | **10.19%** |
| TIR |  |  |  |  |  |  |  |  |  |
| CACT<br>A | 2.17% | 2.07% | 1.88% | 2.18% | 2.00% | 1.65% | 2.16% | 1.51% | **1.95%** |
| Mutato<br>r | 5.90% | 5.85% | 2.53% | 5.73% | 2.76% | 2.79% | 2.28% | 2.76% | **3.83%** |
| PIF_H<br>arbing<br>er | 0.69% | 0.83% | 0.74% | 0.83% | 0.84% | 0.68% | 0.73% | 0.79% | **0.77%** |
| Tc1_M<br>ariner | 2.49% | 2.39% | 2.78% | 2.51% | 2.71% | 2.65% | 2.85% | 2.58% | **2.62%** |
| hAT | 1.68% | 1.72% | 1.75% | 1.75% | 1.76% | 1.71% | 1.79% | 1.69% | **1.73%** |
| nonTI<br>R |  |  |  |  |  |  |  |  |  |
| helitro<br>n | 5.45% | 5.08% | 4.97% | 4.96% | 4.51% | 4.86% | 4.24% | 5.60% | **4.96%** |
| Total | **66.38%** | **65.96%** | **68.35%** | **66.43%** | **67.70%** | **61.90%** | **68.67%** | **69.13%** | **66.82%** |
| % of estimated genome<br>size (1.45Gb) |  |  |  |  |  |  |  |  |  |

#### Iso-Seq library preparation and sequencing

Iso-Seq libraries were produced from 5-6 tissues at different developmental stages for the Regalona and 03-21-072RM genotypes (root, shoot, apical meristem, leaves, and flowers for both genotypes, and developing seeds for Regalona only), in order to support the gene annotation of one lowland (Regalona) and one highland (03-21-072RM) reference genome assembly, respectively. Plants issued from the same seed stock as for the genome sequencing were sown and grown in the same soil, temperature and daylength conditions as described in the DNA extraction section. Each sample was made of tissues collected from three plants. Cleaned tissues washed in pure water were flash frozen in liquid nitrogen and kept at −80°C until processing. RNA extraction was done using RNeasy Mini Kit, Qiagen, Cat. No. 74104, then DNAse treatment was performed using Invitrogen™ DNA-free™ DNA Removal Kit, Catalog No. AM1906. The concentration and purity of the RNA samples were measured on Thermo Scientific™ NanoDrop™ 2000 Spectrophotometer, and RNA integrity was assessed on the Agilent 2100 Bioanalyzer system. Only samples with RNA-integrity (RIN) value above 6 were retained for sequencing. Library preparation following the PacBio Iso-Seq^TM^ Express Template Preparation, and sequencing on Sequel II System was performed at Novogene Co., Ltd.

#### Bionano optical map

##### Genome assembly

For each of the eight quinoa accessions, PacBio HiFi reads were assembled using hifiasm^44^ v16.1 with default parameters to generate primary contig assemblies, and hybrid scaffolds were then generated by combining contig assemblies and optical maps using the hybridScaffold pipeline (https://bionano.com/wp-content/uploads/2023/01/30073-Bionano-Solve-Theory-of-Operation-Hybrid-Scaffold.pdf; Bionano Solve version 3.7). Hybrid scaffolds for each quinoa accession were compared to the quinoa reference QQ74-V2^27^ using a dotplot produced by chromeister^45^. For the accession 03-21-072RM, the dotplot comparison (Fig. 2) showed that the 18 longest (with an average length of 71.2 Mb) hybrid scaffolds out of 21 (with 0.2 Mb average length for the three remaining scaffolds) represented the chromosome-scale assembly, and the orientation and ordering of each pseudomolecule were assigned according to the quinoa reference genome QQ74-V2. Thus, the 03-21-072RM assembly was used as reference to guide the scaffolding of the other quinoa accessions to chromosome-scale level using RagTag^46^ v2.1.0 with the following parameters: -C -r -u. LTR Assembly Index (LAI) was determined with LTR_retriever^47,48^ v2.8.7, and BUSCO^49^ v5.4.5 scores were computed with the eudicots_odb10 lineage dataset to assess the completeness of each genome.

**Fig. 2:**
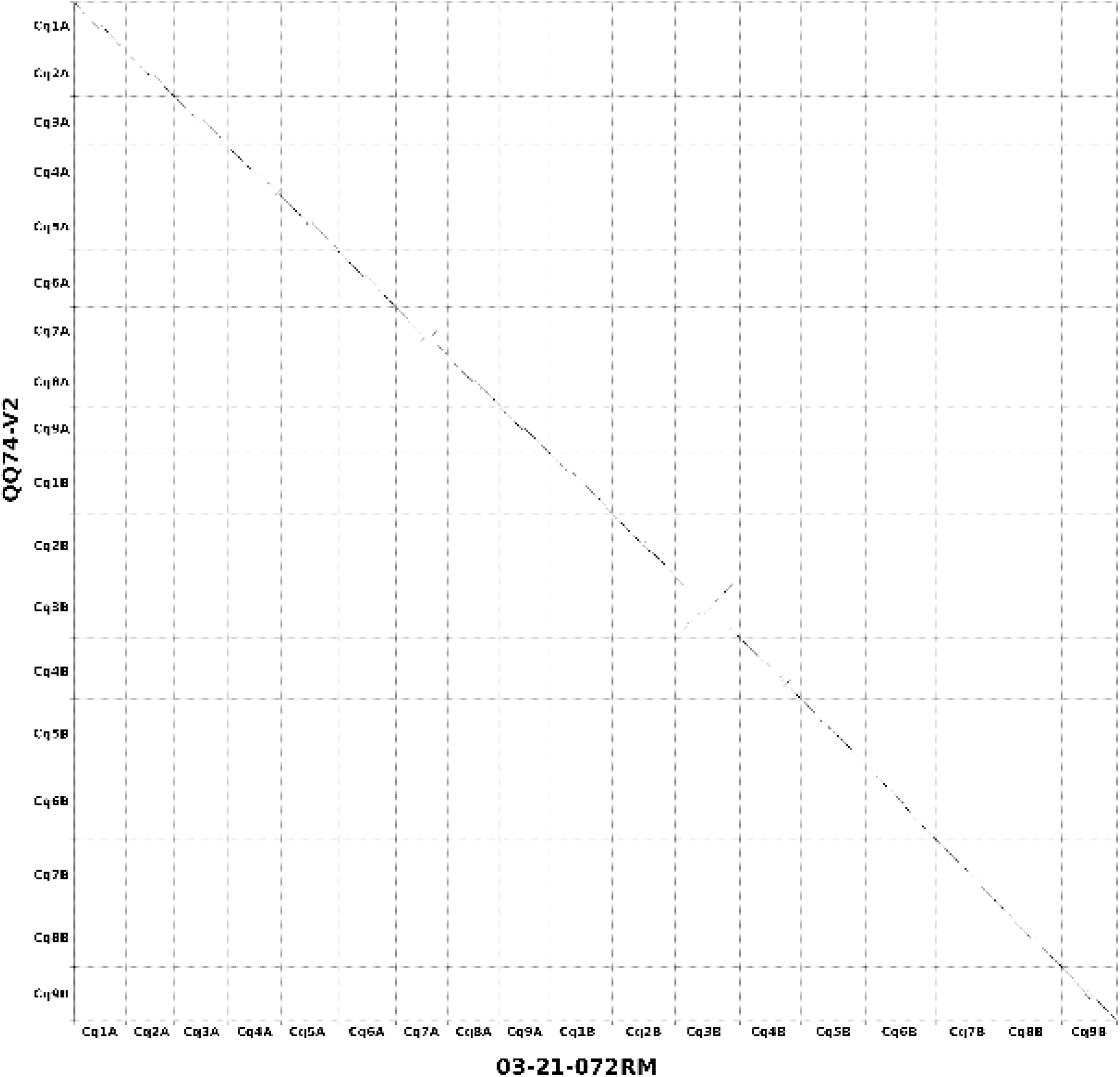
Dotplot comparison of the 18 longest hybrid scaffolds of the quinoa accession 03-21-072RM (x-axis) against the quinoa reference genome assembly QQ74-V2 (y-axis).

### Genome analysis

#### Repeat analysis

We performed the annotation, filtering, and consolidation of the repeat elements across the 8 quinoa genomes using default parameters of panEDTA^50^ with the REPET^51,52^ curated library provided for QQ74-V2^27^ as input. On average, 66% of each quinoa genome is classified as repetitive sequences (Table 2).

#### Gene model prediction

For Regalona and 03-21-072RM accessions, the Circular Consensus Sequencing (CCS) application v6.4.0 (https://github.com/PacificBiosciences/ccs) using the continuous long reads subread dataset and default parameters was used to produce HiFi reads. Iso-Seq tool v4.0.0 (https://github.com/PacificBiosciences/IsoSeq) was used to trim primers and polyadenylation (polyA) tails and produce *de novo* Full Length Non Chimeric (FLNC) transcripts for downstream mapping and annotation.

Then, the FLNC transcripts from the six and five developmental stages were mapped to their corresponding reference assembly using minimap2^53^ v2.17-r941 (parameter: -ax splice -uf – secondary=no -C5 -O6,24 -B4), and the redundant isoforms were further collapsed into transcript loci using cDNA_Cupcake (http://github.com/Magdoll/cDNA_Cupcake; parameter: --dun-merge-5-shorter -C 0.95). All the gff3 files from each sample were merged into a single gtf file for both Regalona and 03-21-072RM accessions using StringTie^54^ v2.1.7. The Transdecoder.LongOrfs script (https://github.com/TransDecoder/TransDecoder/blob/master/TransDecoder.LongOrfs) was used to identify open reading frames (ORF) of at least 100 amino acids from the merged gtf file. The predicted protein sequences were compared to the UniProt (2021_03) and Pfam35 databases using BLASTP^55^ (parameters: -max_target_seqs 1 -outfmt 6 -evalue 1e−5) and hmmer356 v3.3.2 (parameters: hmmsearch -E 1e-10). The Transdecoder.Predict script (https://github.com/TransDecoder/TransDecoder/blob/master/TransDecoder.Predict) was used with the BLASTP and hmmer results to select the best translation per transcript. Finally, the annotation gff3 file was computed using the perl script “cdna_alignment_orf_to_genome_orf.pl” provided in the Transdecoder package.

In addition, a lifting approach using liftoff^57^ v1.6.3 was combined with the genome-guided approach to predict gene models for Regalona and 03-21-072RM accessions. For the gene lifting, the annotations of QQ74-V2 and *Beta vulgaris* cv EL10_v1 were independently transferred using liftoff^57^ (parameters : -a 0.9 -s 0.9 -copies -exclude_partial -polish).

All the output gff files from the lifting and genome-guided approaches were merged into a single file using AGAT (https://github.com/NBISweden/AGAT; perl script “<u>agat_sp_merge_annotations.pl</u>”). The merged file was then post-processed using gffread tool v0.11.7 (parameters: --keep-genes -N -J) to retain transcripts with a start and stop codon and to discard transcripts containing premature stop codons and/or introns with non-canonical splice sites. In total, 46,730 and 43,733 genes were predicted for Regalona and 03-21-072RM, respectively.

For the six others genomes (CHEN90, CHEN199, D10126, D12282, Javi, and PI614919), a gene lifting approach using QQ74-V2, Regalona and 03-21-072RM gene models as reference was performed and merged into a comprehensive gff3 file using the same methods described above (using AGAT (https://github.com/NBISweden/AGAT) and gffread^58^ v0.11.7). Between 43,733 and 48,564 gene models were predicted for the eight new quinoa genomes, and the BUSCO^49^ score was calculated with the protein mode (Table 4). Finally, the putative functional annotations were assigned using a protein comparison against the UniProt database (2021_03) using DIAMOND^59,60^ (parameters: -f 6 -k1 -e 1e-6). PFAM domain signatures and GO terms were assigned using InterproScan^61^ v5.55-88.039.

**Table 4:**
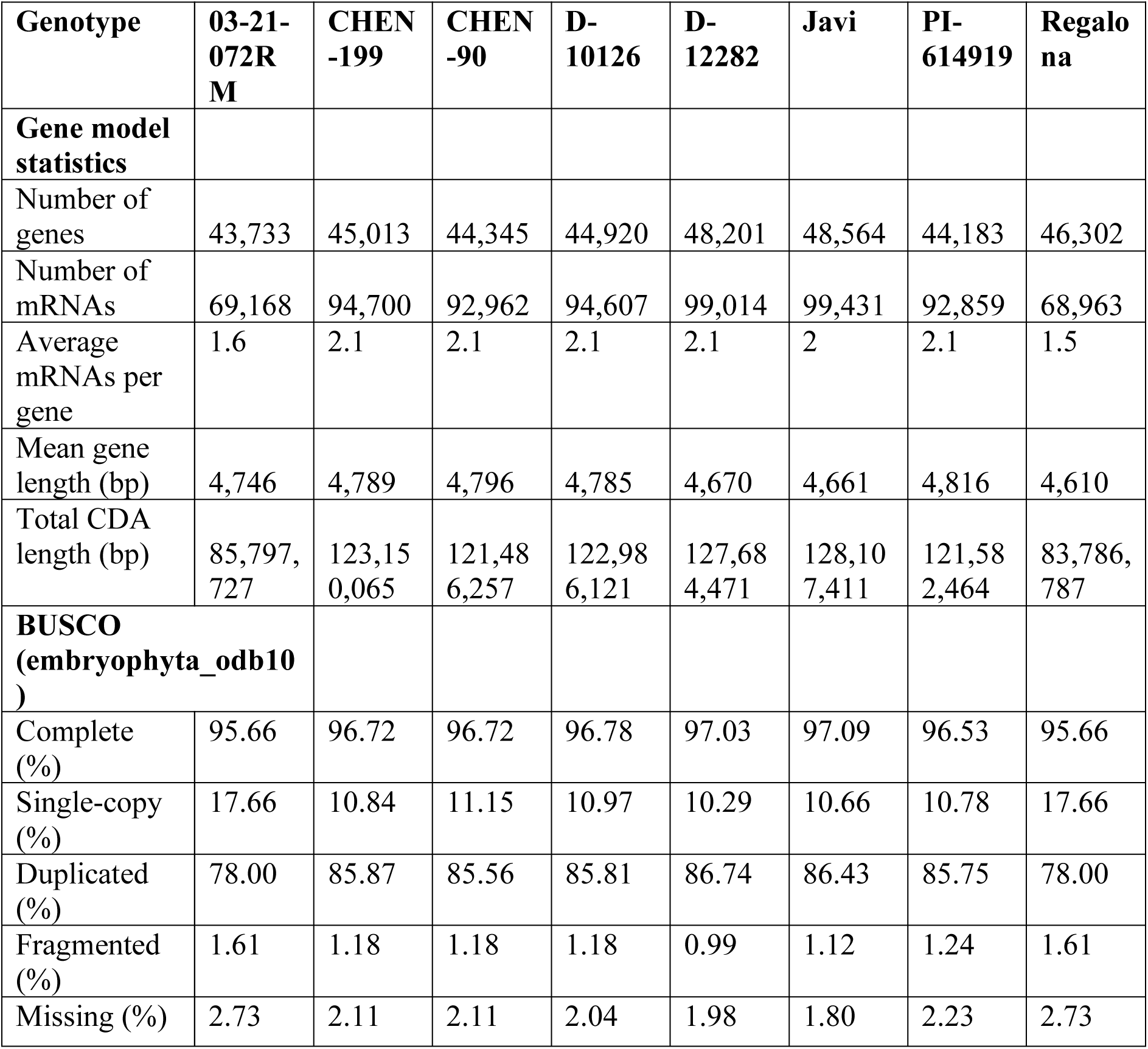
Gene annotation.

| **Genotype** | **03-21-072RM** | **CHEN-199** | **CHEN-90** | **D-10126** | **D-12282** | **Javi** | **PI-614919** | **Regalona** |
| --- | --- | --- | --- | --- | --- | --- | --- | --- |
| **Gene model statistics** |  |  |  |  |  |  |  |  |
| Number of genes | 43,733 | 45,013 | 44,345 | 44,920 | 48,201 | 48,564 | 44,183 | 46,302 |
| Number of mRNAs | 69,168 | 94,700 | 92,962 | 94,607 | 99,014 | 99,431 | 92,859 | 68,963 |
| Average mRNAs per gene | 1.6 | 2.1 | 2.1 | 2.1 | 2.1 | 2 | 2.1 | 1.5 |
| Mean gene length (bp) | 4,746 | 4,789 | 4,796 | 4,785 | 4,670 | 4,661 | 4,816 | 4,610 |
| Total CDA length (bp) | 85,797,727 | 123,150,065 | 121,486,257 | 122,986,121 | 127,684,471 | 128,107,411 | 121,582,464 | 83,786,787 |
| **BUSCO (embryophyta_odb10)** |  |  |  |  |  |  |  |  |
| Complete (%) | 95.66 | 96.72 | 96.72 | 96.78 | 97.03 | 97.09 | 96.53 | 95.66 |
| Single-copy (%) | 17.66 | 10.84 | 11.15 | 10.97 | 10.29 | 10.66 | 10.78 | 17.66 |
| Duplicated (%) | 78.00 | 85.87 | 85.56 | 85.81 | 86.74 | 86.43 | 85.75 | 78.00 |
| Fragmented (%) | 1.61 | 1.18 | 1.18 | 1.18 | 0.99 | 1.12 | 1.24 | 1.61 |
| Missing (%) | 2.73 | 2.11 | 2.11 | 2.04 | 1.98 | 1.80 | 2.23 | 2.73 |

#### Genome visualization

The eight quinoa genomes of were uploaded into the Persephone^®^ multi-genome browser (https://web.persephonesoft.com/). The data tracks available are the DNA sequence and gene model prediction. Among other functions provided by the platform, a BLAST^36^ search and synteny analysis within all quinoa genomes is also available (Fig. 3).

**Fig 3:**
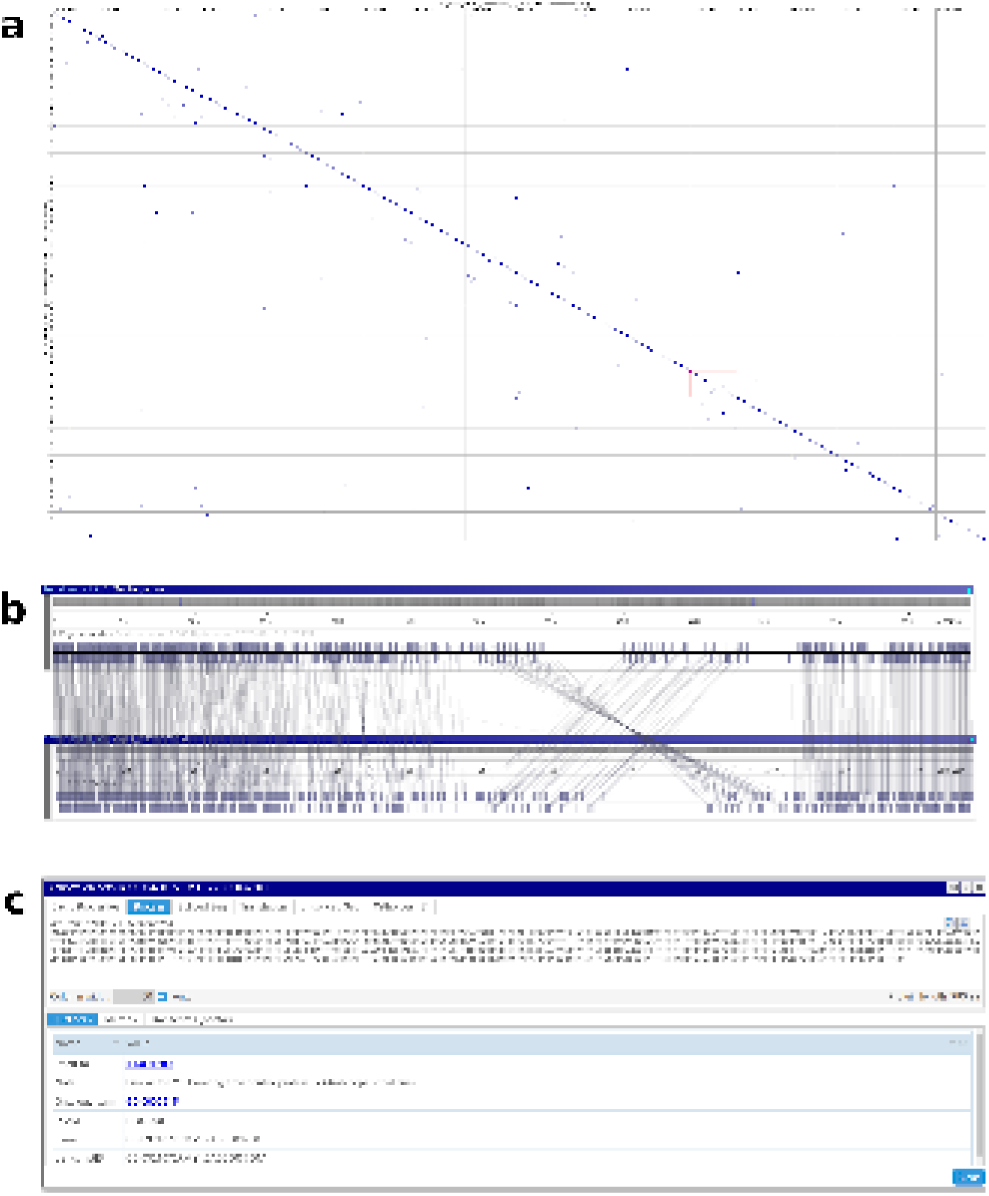
Visualization of quinoa genomes using Persephone^®^ genome browser. Example of visualization of the synteny between a) all chromosomes and b) Cq7A chromosome of Regalona and 03-21-072RM genomes, and c) gene and related sequence search function.

#### Structural variation

The collinearity and major structural variations (inversions, duplications, and translocations) between QQ74 and the eight new quinoa genomes were assessed using minimap2^53^ v2.26-r1175 (parameters: -ax asm5 --eqx) and SyRI^62^ v1.6.3 with defaults parameters. The results were visualized with plotsr^63^ v1.1.1 (Fig. 4).

**Fig 4:**
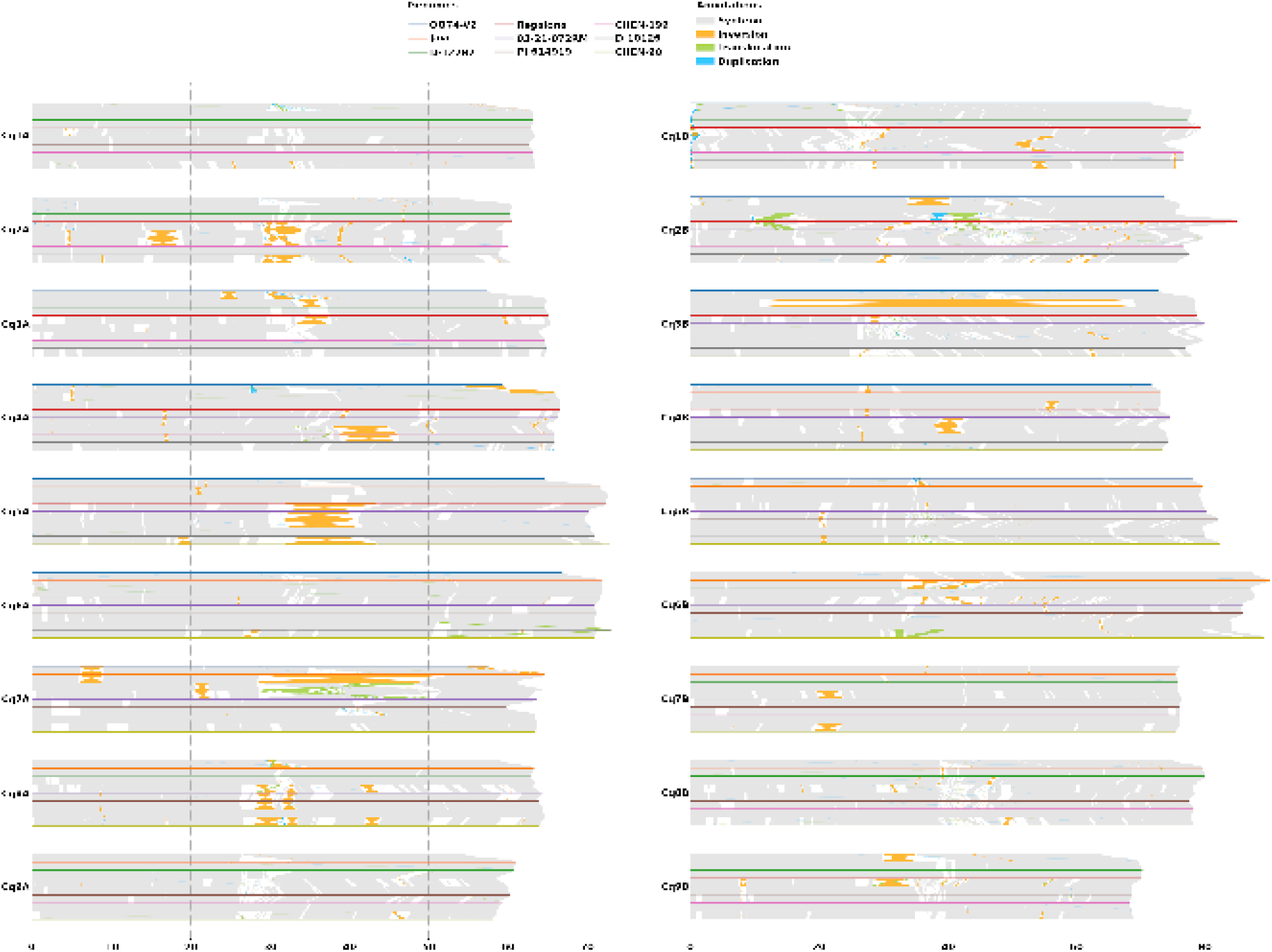
Collinearity and structural rearrangements between QQ74-V2 and the eight quinoa genome assemblies identified using SyRI and visualized with plotsr. On the left and right panels are shown alignments for each chromosome of the A and B sub-genomes, respectively. For each chromosome, the different genomes are ordered per similarity from top to bottom as follows: QQ74-V2, Javi, D-12282, and Regalona for the lowland accessions; 03-21-072RM, PI-614919, CHEN-199, D-10126, and CHEN-90 for the highland accessions. Positions along the chromosomes are given on the x-axes in Mb. Syntenic regions between genomes are shown in grey, inversions in yellow, translocations in green, and duplications in blue.

All chromosomes from both quinoa sub-genomes present at least one type of chromosomal rearrangement. The previously identified Cq3B large pericentric inversion (In(3B)(11136405::63361214))^27^ is shared between the QQ74 and Javi genotypes and is the largest chromosomal rearrangement detected among all genotypes. Large chromosomal inversions on Cq5A and Cq7A are also present in multiple genomes. The pseudomolecules for QQ74-V2 are shorter for most of the chromosomes in comparison to the eight new reference sequences, with notable collapse of sequences in the centromeric and telomeric regions. This can be explained by the different methods of sequencing (CLR PacBio reads) and assembly software used for the QQ74 genome assembly which resulted in lower sequence completeness and contiguity in comparison to the new genomes produced with PacBio HiFi sequencing and assembly technology supported by the newest Bionano optical maps for scaffolding (Table 2).

### Data Records

The HiFi and Iso-Seq reads and the final chromosome assemblies were deposited in the Sequence Read Archive at NCBI under BioProject number PRJNA1018548. The raw Bionano optical maps were deposited under EMBL project number PRJEB66274.

The quinoa assemblies, gene model prediction and the repeat annotations are available on DRYAD Digital Repository (doi:10.5061/dryad.zkh1893jj).

### Technical validation

#### Assessment of genome assembly and annotation

The average BUSCO^49^ v5.4.5 (using embryophyta_odb10) score for the eight quinoa genomes is 98.1% at the genome level, indicating a high completeness of the assemblies. In comparison, the BUSCO^49^ score for the previous quinoa genome QQ74-V2 is 97.9%. The quality of the eight genome assemblies was also assessed with Merqury based on the PacBio HiFi reads using 19-mers. The QV (consensus quality value) scores are in the range of 64.7 and 69 for CHEN199 and PI614919, respectively. The k-mer completeness scores were between 98.25% and 99.03 for Regalona and D10126 genomes, respectively (Table 2).

We validated the concordance of the assembly by re-mapping the optical map onto the pseudomolecules of each of the eight quinoa genomes. Each pseudomolecule was visually inspected using Bionano Access software, and no major discrepancies were found. Telomeric repeats (TTTAGGG)n ^27^ were screened using tidk^64^ v0.2.31 (parameter: find -c Caryophyllales) (https://github.com/tolkit/telomeric-identifier). Telomeric repeats are present at the extremities of 35 out of 36 pseudomolecules for the assemblies of 03-21-072RM, CHEN90, D10126 and Regalona. Interestingly, telomeric repeat sequences are absent from the short arm of Cq1B in all genome assemblies. Telomeric repeat sequences are missing for one additional chromosome for CHEN199 (Cq2BL), D12282 (Cq2BS) and PI614919 (Cq8AS), while Javi has missing telomere for two additional chromosomes (Cq2AS and Cq7AS).

Completeness of the gene model prediction was evaluated using BUSCO^49^ v5.4.5 (using embryophyta_odb10) and produced a score between 95.6% for Regalona and 97.1% for Javi.

### Code availability

All software and pipelines were executed according to the manual and protocol of published tools. No custom code was generated for these analyses.

## Acknowledgments

We would like to thank Ingrid von Baer for enabling the use of Regalona quinoa in this study, and the public release of all related data to the scientific community. We thank the KAUST Bioscience Core Laboratory for sequencing support, the KAUST Plant Growth Facility Core Lab for greenhouse support, and the KAUST Supercomputing Facilities (https://www.hpc.kaust.edu.sa/) for providing computing resources. This research was supported by the King Abdullah University of Science and Technology (KAUST).

## Author contribution

E.R. and M.Tester conceived the project and planned the analyses. E.R., G.F., C.S., D.E.J., E.N.J., P.J.M., and M.Tester contributed to the germplasm selection. E.R. performed the gDNA extractions, sequence assemblies and validations. J.C. and E.H. performed the genome size estimation analyses. N.R., I.D., and S.C. produced the optical maps and hybrid scaffold assemblies. N.S. performed the RNA extractions. M.A. performed the annotations and variants identification. M.T. and M.K. managed the visualization platform. E.R., M.A., and M.Tester wrote the initial manuscript with input from all authors. All authors have read and approved the final manuscript.

## Competing interests

The authors declare no competing interests.

## Additional information

Correspondence and requests should be addressed to E.R or M. Tester.

## Notes

### Competing Interest Statement

The authors have declared no competing interest.

